# ComPotts: Optimal alignment of coevolutionary models for protein sequences

**DOI:** 10.1101/2020.06.12.147702

**Authors:** Hugo Talibart, François Coste

## Abstract

To assign structural and functional annotations to the ever increasing amount of sequenced proteins, the main approach relies on sequence-based homology search methods, e.g. BLAST or the current state-of-the-art methods based on profile Hidden Markov Models (pHMMs), which rely on significant alignments of query sequences to annotated proteins or protein families. While powerful, these approaches do not take coevolution between residues into account. Taking advantage of recent advances in the field of contact prediction, we propose here to represent proteins by Potts models, which model direct couplings between positions in addition to positional composition. Due to the presence of non-local dependencies, aligning two Potts models is computationally hard. To tackle this task, we introduce an Integer Linear Programming formulation of the problem and present ComPotts, an implementation able to compute the optimal alignment of two Potts models representing proteins in tractable time. A first experimentation on 59 low sequence identity pairwise alignments, extracted from 3 reference alignments from sisyphus and BaliBase3 databases, shows that ComPotts finds better alignments than the other tested methods in the majority of these cases.

## 1 Introduction

Thanks to sequencing technologies, the number of available protein sequences has considerably increased in the past years, but their functional and structural annotation remains a bottleneck. This task is thus classically performed *in silico* by scoring the alignment of new sequences to well-annotated homologs. One of the best-known method is BLAST[1], which performs pairwise sequence alignments. The main tools for homology search use now Profile Hidden Markov Models (pHMMs), which model position-specific composition, insertion and deletion probabilities of families of homologous proteins. Two well-known software packages using pHMMs are widely used today: HMMER[2] aligns sequences to pHMMs and HH-suite[3] takes it further by aligning pHMMs to pHMMs.

Despite their solid performance, pHMMs are innerly limited by their positional nature. Yet, it is well-known that residues that are distant in the sequence can interact and co-evolve, e.g. due to their spatial proximity, resulting in correlated positions (see for instance [4]).

There have been a few attempts to make use of long-distance information. Menke, Berger and Cowen introduced a Markov Random Field (MRF) approach where MRFs generalize pHMMs by allowing dependencies between paired residues in *β*-strands to recognize proteins that fold into *β*-structural motifs[5]. Their MRFs are trained on multiple structure alignments. Simplified models[6] and heuristics[7] have been proposed to speed up the process. While these methods outperform HMMER[2] in propeller fold prediction, they are limited to sequence-MRF alignment on *β*-strand motifs with available structures. Xu et al.[8] proposed a more general method, MRFalign, which performs MRF-MRF alignments using probabilities estimated by neural networks from amino acid frequencies and mutual information. Unlike SMURF, MRFalign allows dependencies between all positions and MRFs are built on multiple sequence alignments. MRFalign showed better alignment precision and recall than HHalign and HMMER on a dataset of 60K non-redundant SCOP70 protein pairs with less than 40% identity with respect to reference structural alignments made by DeepAlign[9], showing the potential of using long-distance information in protein sequence alignment.

Meanwhile, another type of MRF led to a breakthrough in the field of contact prediction[10]: the Potts model. This model was brought forward by Direct Coupling Analysis[11], a statistical method to extract direct correlations from multiple sequence alignments. Once inferred on a MSA, a Potts model’s nodes represent positional conservation, and its edges represent direct couplings between positions in the MSA. Unlike mutual information which also captures indirect correlations between positions, Potts models are global models capturing the collective effects of entire networks of correlations through their coupling parameters[12], thus tackling indirect effects and making them a relevant means of predicting interactions between residues. Beyond contact prediction, the positional and the direct coupling information captured by Potts model’s parameters might also be valuable in the context of protein homology search. The idea of using Potts models for this purpose was proposed last year at the same workshop by Muntoni and Weigt[13], who propose to align sequences to Potts models, and by us[14] with the introduction of ComPotts, our method to align Potts models to Potts models.

In this paper, we fully describe ComPotts and focus on its performances in terms of alignment quality. In the following sections, we explain our choices for Potts model inference and we describe our method for aligning them, which builds on the work of Wohlers, Andonov, Malod-Dognin and Klau[15,16,17] to propose an Integer Linear Programming formulation for this problem, with an adequate scoring function. We assess the quality of ComPotts’ alignments with respect to 59 reference pairwise alignments extracted from sisyphus[18] and BaliBase3[19] databases. On these first experiments, computation time was tractable and ComPotts found better alignments than its main competitors: BLAST, HHalign (which is HHblits’ alignment method) and MRFalign.

## 2 Methods

In this section, we describe our approach to align two Potts models. We start with a short summary of Potts models notations and then we explain the choices we made for the inference of Potts models. Then, we introduce our formulation of the alignment problem as an Integer Linear Programming problem, using notations from [20].

### 2.1 Inference of Potts models

Potts models are discrete instances of pairwise Markov Random Fields which originate from statistical physics. They generalize Ising models by describing interacting spins on a crystalline lattice with a finite alphabet. In the paper introducing Direct Coupling Analysis[11], Weigt et al. came up with the idea of applying them to proteins: inferred on a multiple sequence alignment, a Potts Model could then be used to predict contacts between residues.

A Potts model on protein sequences can be defined as follows:

Let *S* be a multiple sequence alignment (MSA) of length *L* over an alphabet Σ of length *q* (here we use the amino acid alphabet, which is of length *q* = 20). A Potts model for *S* is a statistical model defining a probability distribution over the set Σ^*L*^ of all sequences of length *L* which complies to the maximum-entropy principle and whose single and double marginal probabilities are the empirical frequencies of the MSA. Formally, denoting *f*_*i*_(*a*) the frequency of letter *a* at position *i* in the MSA *S* and *f*_*ij*_(*a, b*) the frequency of *a* and *b* together at positions *i* and *j* in *S*, a Potts model for *S* satisfies:

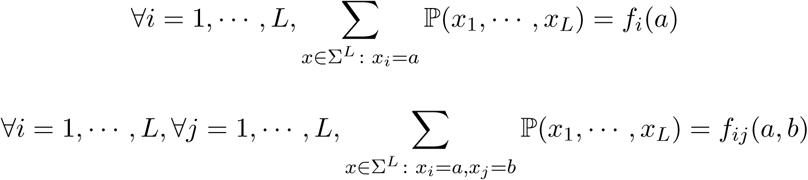

and defines a Boltzmann distribution on Σ^*L*^ with:

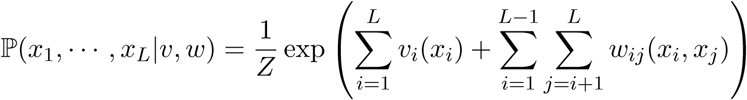

where:

— *Z* is a normalization constant : 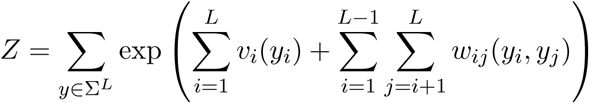
— {*v*_*i*_}_*i*=1,, *L*_ are positional parameters termed *fields*. Each *v*_*i*_ is a real vector of length *q* where *v*_*i*_(*a*) is a weight related to the propensity of letter *a* to be found at position *i*.
— {*w*_*ij*_}_*i,j*=1*…,L*_ are *pairwise couplings*. Each *w*_*ij*_ is a *q* × *q* real weight matrix where *w*_*ij*_(*a, b*) quantifies the tendency of the letters *a* and *b* to co-occur at positions *i* and *j*.

The value 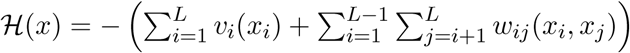 is called Hamiltonian.

In theory, one could infer a Potts model from a MSA *S* by likelihood maximization, i.e. by finding the positional parameters *v* and coupling parameters *w* that maximize ℙ(*S*|*v, w*). In practice, however, this would require the computation of the normalization constant *Z* at each step, which is computationally intractable. Among the several approximate inference methods that have been proposed [21,22,23,24,12], we opted for pseudo-likelihood maximization since it was proven to be a consistent estimator in the limit of infinite data [25,26] within reasonable time. Furthermore, since our goal is to align Potts models, we need the inferrence to be geared towards similar models for similar MSAs, which is not what inference methods were initially designed for. In an effort towards inferring canonical Potts models, we chose to use CCMpredPy[27], a recent Python-based version of CCMpred[28] which, instead of using the standard *L*_2_ regularization prior 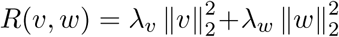, uses a smarter prior on 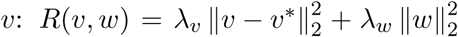 where *v*\* obeys 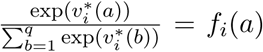 which yields the correct probability model if no columns are coupled, i.e.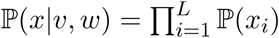. Our intuition is that positional parameters should explain the MSA as much as possible and only necessary couplings should be added.

### 2.2 Alignment of Potts models

We introduce here our method for aligning two Potts models. We start by describing the function we designed to score a given alignment, then we add the constraints that make the alignment proper by embedding it in an Integer Linear Programming formulation, following Wohlers et al.[17], allowing us to use their efficient solver for the optimization.

#### 2.2.1 Scoring an alignment

We want the alignment score of two Potts models *A* and *B* to maximize the similarity between aligned fields and aligned couplings.

Formally, we want to find the binary variables *x*_*ik*_ and *y*_*ikjl*_, where *x*_*ik*_ = 1 iff node *i* of Potts model *A* is aligned with node *k* of Potts Model *B* and *y*_*ikjl*_ = 1 iff edge (*i, j*) of Potts model *A* is aligned with edge (*k, l*) of Potts model *B*, such that: 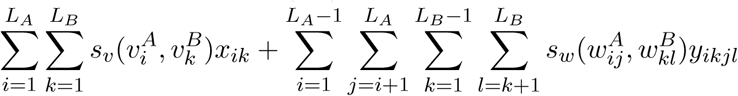 is maximized, where 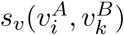 and 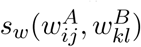 are similarity scores between positional parameters 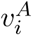 and 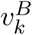 and coupling parameters 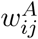 and 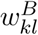.

To score the similarity 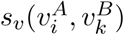 between positional parameters 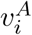 and 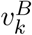 we use the scalar product :

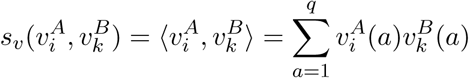

And to score the similarity 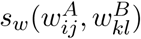 between coupling parameters 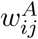 and 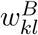 we use the Frobenius inner product, which is the natural extension of scalar product to matrices :

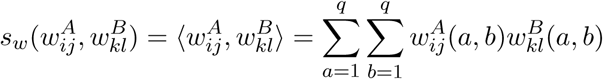

This scoring function can be seen as a natural extension of the opposite of the Hamiltonian of a sequence *x*, since 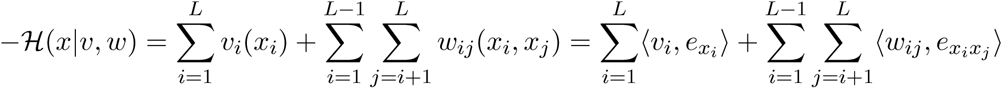 where : — 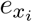 is the vector defined by ∀*a* ∈ [1..*q*], 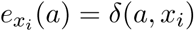

— 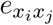 is the matrix defined by ∀(*a, b*) ∈ [1..*q*]^2^, 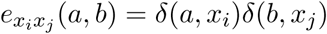

#### 2.2.2 Optimizing the score with respect to constraints

Naturally, the scoring function should be maximized with respect to constraints ensuring that the alignment is proper. In that perspective, we build on the work of Wohlers et al.[17], initially dedicated to protein structure alignment, to propose an Integer Linear Programming formulation for the Potts model alignment problem.

We remind first the constraints for a proper alignment following [20].

First, we need the definition of *alignment graph*. For *A* and *B* two Potts Models of lengths *L*_*A*_ and *L*_*B*_, the alignment graph *G* is a *L*_*A*_ × *L*_*B*_ grid graph where rows (from bottom to top) represent the nodes of *A* and columns (from left to right) represent the nodes of *B*. A node *i*.*k* in the alignment graph indicates the alignment of node *i* from Potts model *A* and node *k* from Potts model *B*.

Every proper alignment of two Potts model is described by a *strictly increasing path* in this alignment graph, which is defined as a subset {*i*_1_.*k*_1_, *i*_2_.*k*_2_, …, *i*_*n*_.*k*_*n*_} of alignment graph nodes that can be ordered such that each node is strictly larger than the previous one, i.e. *i*_1_ < *i*_2_ < … < *i*_*n*_ and *k*_1_ < *k*_2_ < … < *k*_*n*_.

To specify the constraints of the ILP, they defined sets of mutually contradicting nodes, called *decreasing paths*. A decreasing path is a set {*i*_1_.*k*_1_, *i*_2_.*k*_2_, …, *i*_*n*_.*k*_*n*_} of alignment graph nodes for which *i*_1_ ≥ *i*_2_ ≥ … ≥ *i*_*n*_ and *k*_1_ ≤ *k*_2_ ≤ … ≤ *k*_*n*_ holds. The set of all decreasing paths is denoted *C*.

We also give notations for the left and right neighborhood of a node : let *i*.*k* be a node in the alignment graph and 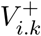 (resp. 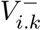) denote the set of couples that are strictly larger (resp. smaller) than *i*.*k*, e.g.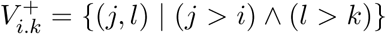 and 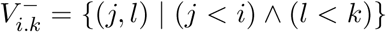 and let 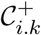 (resp.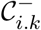) denote the set of all decreasing paths in 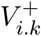 (resp. 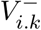).

Given the notations above, with *A* (resp. *B*) a Potts Model of length *L*_*A*_ (resp. *L*_*B*_) with parameters *v*^*A*^ and *w*^*A*^ (resp *v*^*B*^ and *w*^*B*^), aligning *A* and *B* can be formulated as the following Integer Linear Programming problem:

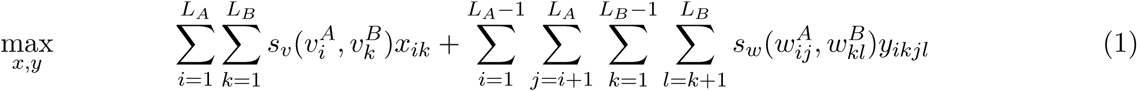

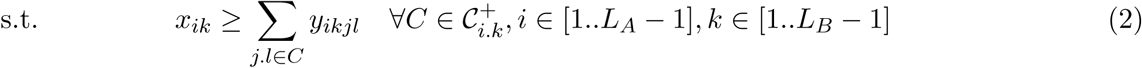

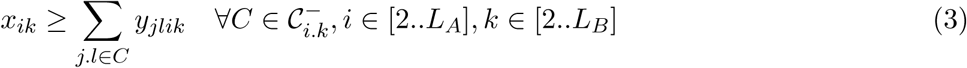

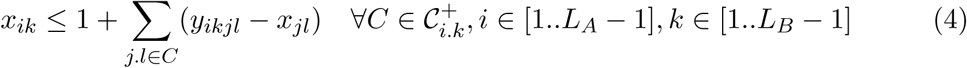

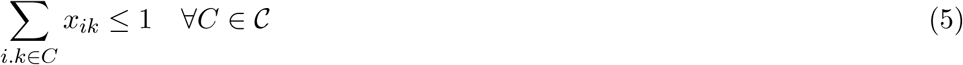

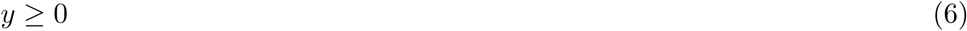

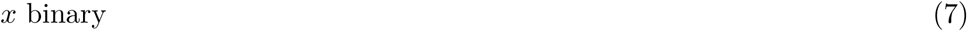

As in [17], an affine gap cost function can be added to the score function to account for insertions and deletions in the sequences.

## 3 Results

We implemented this ILP formulation in a program, ComPotts, embedding the solver from [17]. We assessed the performances of ComPotts in terms of alignment precision and recall with respect to a set of 59 pairwise reference alignments. For each sequence, a Potts model was inferred on a multiple sequence alignment of close homologs retrieved by HHblits.

### 3.1 Data

We extracted 59 reference pairwise sequence alignments from 3 reference multiple sequence alignments from sisyphus[18] and BaliBase3[19] with a particularly low sequence identity. To focus on sequences with coevolution information, we considered only sequences with at least 1000 close homologs (see next section). We also discarded sequences with more than 200 amino acids for memory considerations with respect to CCMpredPy. Reference alignments are summed up in table 3.1

**Tab. 1.**
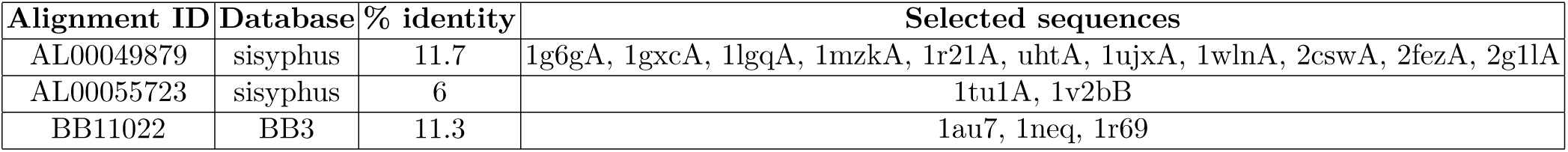
Reference multiple alignments used in our experiment and selected sequences extracted.

### 3.2 Alignment experiment

#### 3.2.1 Potts model inference

For each sequence, we built a MSA of its close homologs by running HHblits[3] v3.0.3 on Uniclust30[29] (version 08/2018) with parameters recommended in [30] for CCMpred: `-maxfilt 100000 -realign_max 100000 -all -B 100000 -Z 100000 -n 3 -e 0.001` which we then filtered at 80% identity using HHfilter, and took the first 1000 sequences. If the MSA had less than 1000 sequences we removed it from the experiment. This MSA was then used to train a Potts model with CCMpredPy using default parameters except for the *w* regularization factor coefficient (we set it to 30, which we empirically found to result in 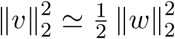, in other words making *v* and *w* scores commensurable) and the uniform pseudo-counts count on the *v* (we set it to 1000 to have as many pseudo-counts as the number of sequences in the alignment in order to enhance stronger conservation signal and limit excessive similarity scores due to missing the same rare residues).

#### 3.2.2 Potts model alignment

We ran ComPotts with a gap open cost of 8 and no gap extend cost, which we found empirically to yield the best alignments in previous experiments. To speed up the computations, we decided to stop the optimization when 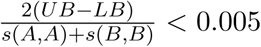 with *UB* and *LB* the current upper and lower bounds of the Lagrangian optimization, since we realized in preliminary experiments that in practice it gave the same alignments as the optimal ones in significantly less time.

### 3.3 Alignment quality assessment

We compared each resulting alignment with the reference pairwise alignment extracted from the multiple sequence alignment, considering the alignment precision with the Modeler score[31] 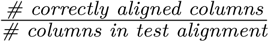 and the alignment recall with the TC score[32] 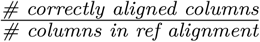, computed using Edgar’s qscore program[33] v2.1.

To compare our results, we ran HHalign v3.0.3 to align HMMs built on the MSAs outputted by HHblits, MRFalign v0.90 to align MRFs it built from the sequences, both with default options, and BLASTp v2.9.0+ without E-value cutoff. As a control, we also ran Matt v1.00 on the corresponding PDB structures. Results are summarized in figure 2. Note that Matt failed to run 3 of the alignments.

**Fig. 1.**
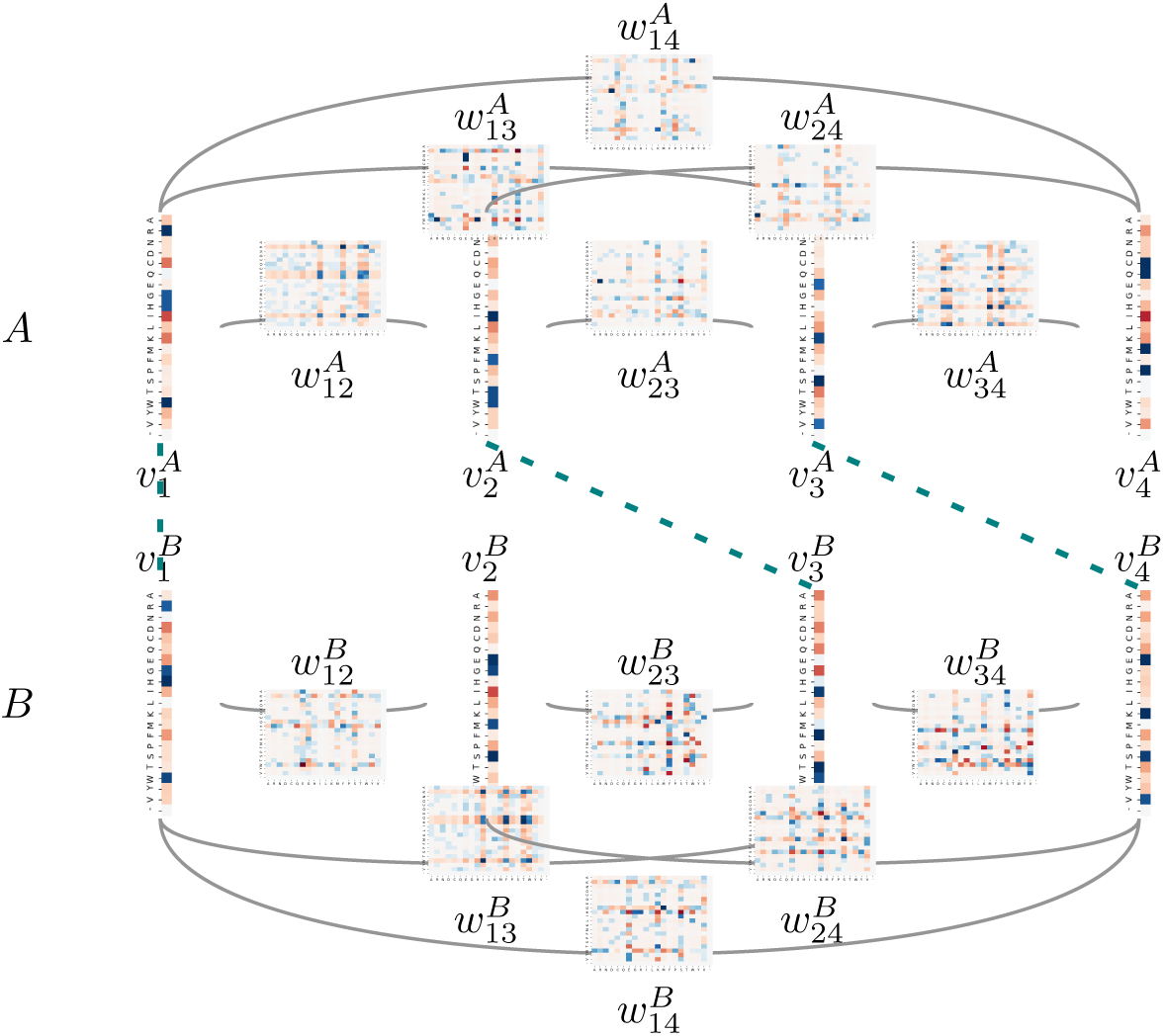
Illustration of the alignment of two Potts models *A* and *B*.

**Fig. 2.**
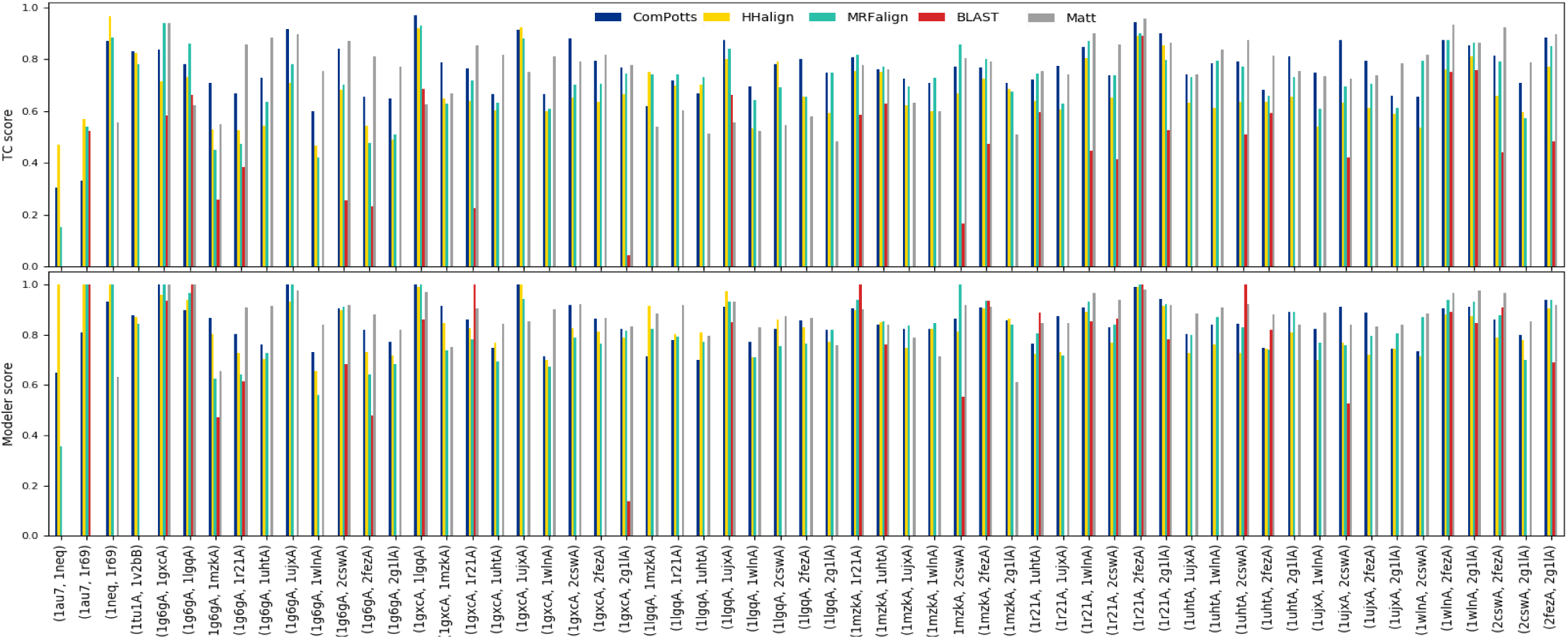
TC score (alignment recall) and Modeler score (alignment precision) for all 59 alignments.

BLAST is unquestionably outperformed by all other tools on this set. 10 out of the 59 sequence pairs could not be aligned (not hit was found) and, on 22 of the alignments it performed, BLAST had both a recall (TC score) and a precision (Modeler score) of 0. Its average TC score is 0.2694 and its average Modeler score is 0.4357, which is about half the average scores of the other methods. BLAST has a better precision on some alignments, most of the time because its alignments are smaller, which results in a rather low recall, except for some alignments which seem to be quite easy for everyone, such as 1r21A and 2fezA.

All methods seem to struggle with the alignment of 1au7 and 1neq: HHalign’s precision skyrockets to 1.0, but at the cost of a recall of 0.47, while ComPotts and MRFalign yield their worst scores, with respective recalls of 0.31 and 0.65 and respective precisions of 0.15 and 0.35.

On average, ComPotts’ alignments have a better recall than all compared tools including Matt with 0.758, versus 0.670 for HHalign, 0.713 for MRFalign, and 0.749 for Matt, outperforming HHalign most of the time – in 52 out of the 59 alignments – and MRFalign in 39 alignments out of the 59, while still having a slightly better precision than all other sequence-based tools with 0.847 while HHalign’s is 0.826 and MRFalign’s is 0.822, outperforming HHalign in 46 alignments out of the 59 and MRFalign in 30 alignments. Matt has the best precision on average with 0.872. Overall, ComPotts has an average *F*1 score 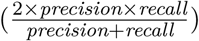 of 0.800, versus 0.740 for HHalign and 0.763 for MRFalign, yielding better alignments than HHalign in 52 cases and better than MRFalign in 39 cases. For asyet-unknown reasons though, our scores for the alignment of 1au7 and 1r69 are remarkably lower than our competitors.

### 3.4 Computation time considerations

We examined the computation times of ComPotts, HHalign and MRFalign, considering only the time they took to align the models and not the time needed to build the models. Not surprisingly, ComPotts is significantly slower than HHalign and MRFalign. This is explained by the fact that HHalign only performs 1D alignment, and MRFalign uses a heuristic to compute the alignment, whereas ComPotts uses an exact algorithm. Aligning two sequences took between 37 seconds (for two models with 75 and 63 positions) and 14.49 minutes (for two models with 144 and 151 positions), with an average of 3.33 minutes on a Debian9 virtual machine with 4 vCPUs and 8GB of RAM, whereas HHalign yields a solution in less than 4 seconds and MRFalign in less than 0.20 seconds. It is worth noting that, although the computation time is significantly higher than its competitors, the solver yields an exact solution in tractable time, even though this problem is NP-complete[34]. In this experiment, the computation time seems to be dominated by the computation of all the *s*_*w*_ scores, which is quadratic in the number of pairs of edges.

## 4 Conclusion

We described ComPotts, our ILP-based method for Potts model-Potts model alignment which can yield the exact solution in tractable time. We reported encouraging results on first experiments where ComPotts often yields better alignments than its two main competitors, HHalign and MRFalign, with respect to a set of 59 low sequence identity reference pairwise alignments. These initial results suggest that direct coupling information can improve protein sequence alignment and might improve sequence-based homology search as well. We still have to see whether the score yielded by ComPotts has more discriminatory power than other methods and enables to better distinguish homologous from non-homologous proteins.

## Acknowledgements

HT is supported by a PhD grant from *Ministère de l’Enseignement Supérieur et de la Recherche* (MESR). We would like to warmly thank Inken Wohlers for providing us with her code, and Mathilde Carpentier for providing a selection of difficult reference alignments and helpful scripts for alignment assessment.

## References

[1] Stephen F Altschul, Warren Gish, Webb Miller, Eugene W Myers, and David J Lipman. Basic local alignment search tool. Journal of molecular biology, 215(3):403–410, 1990.

[2] Sean R. Eddy. Profile hidden markov models. Bioinformatics (Oxford, England), 14(9):755–763, 1998.

[3] Martin Steinegger, Markus Meier, Milot Mirdita, Harald Voehringer, Stephan J Haunsberger, and Johannes Soeding. Hh-suite3 for fast remote homology detection and deep protein annotation. bioRxiv, page 560029, 2019.

[4] Michael Socolich, Steve W Lockless, William P Russ, Heather Lee, Kevin H Gardner, and Rama Ranganathan. Evolutionary information for specifying a protein fold. Nature, 437(7058):512, 2005.

[5] Matt Menke, Bonnie Berger, and Lenore Cowen. Markov random fields reveal an n-terminal double beta-propeller motif as part of a bacterial hybrid two-component sensor system. Proceedings of the National Academy of Sciences, 107(9):4069–4074, 2010.

[6] Noah M Daniels, Raghavendra Hosur, Bonnie Berger, and Lenore J Cowen. Smurflite: combining simplified markov random fields with simulated evolution improves remote homology detection for beta-structural proteins into the twilight zone. Bioinformatics, 28(9):1216–1222, 2012.

[7] Noah M Daniels, Andrew Gallant, Norman Ramsey, and Lenore J Cowen. Mrfy: remote homology detection for beta-structural proteins using markov random fields and stochastic search. IEEE/ACM transactions on computational biology and bioinformatics, 12(1):4–16, 2014.

[8] Jianzhu Ma, Sheng Wang, Zhiyong Wang, and Jinbo Xu. Mrfalign: protein homology detection through alignment of markov random fields. PLoS computational biology, 10(3):e1003500, 2014.

[9] Sheng Wang, Jianzhu Ma, Jian Peng, and Jinbo Xu. Protein structure alignment beyond spatial proximity. Scientific reports, 3:1448, 2013.

[10] Bohdan Monastyrskyy, Daniel D’Andrea, Krzysztof Fidelis, Anna Tramontano, and Andriy Kryshtafovych. New encouraging developments in contact prediction: Assessment of the casp 11 results. Proteins: Structure, Function, and Bioinformatics, 84:131–144, 2016.

[11] Martin Weigt, Robert A White, Hendrik Szurmant, James A Hoch, and Terence Hwa. Identification of direct residue contacts in protein–protein interaction by message passing. Proceedings of the National Academy of Sciences, 106(1):67–72, 2009.

[12] Matteo Figliuzzi, Pierre Barrat-Charlaix, and Martin Weigt. How pairwise coevolutionary models capture the collective residue variability in proteins? Molecular biology and evolution, 35(4):1018–1027, 2018.

[13] Anna Paola Muntoni, Andrea Pagnani, Martin Weigt, and Francesco Zamponi. Using direct coupling analysis for the protein sequences alignment problem. In CECAM 2019 - workshop on Co-evolutionary methods for the prediction and design of protein structure and interactions, 2019.

[14] Hugo Talibart and François Coste. Using residues coevolution to search for protein homologs through alignment of potts models. In CECAM 2019 - workshop on Co-evolutionary methods for the prediction and design of protein structure and interactions, 2019.

[15] Rumen Andonov, Noël Malod-Dognin, and Nicola Yanev. Maximum contact map overlap revisited. Journal of Computational Biology, 18(1):27–41, 2011.

[16] Inken Wohlers, Rumen Andonov, and Gunnar W Klau. Algorithm engineering for optimal alignment of protein structure distance matrices. Optimization Letters, 5(3):421–433, 2011.

[17] Inken Wohlers, Rumen Andonov, and Gunnar W Klau. Dalix: optimal dali protein structure alignment. IEEE/ACM Transactions on Computational Biology and Bioinformatics, 10(1):26–36, 2012.

[18] Antonina Andreeva, Andreas Prlić, Tim JP Hubbard, and Alexey G Murzin. Sisyphus—structural alignments for proteins with non-trivial relationships. Nucleic acids research, 35(suppl 1):D253–D259, 2007.

[19] Julie D Thompson, Patrice Koehl, Raymond Ripp, and Olivier Poch. Balibase 3.0: latest developments of the multiple sequence alignment benchmark. Proteins: Structure, Function, and Bioinformatics, 61(1):127–136, 2005.

[20] Inken Wohlers. Exact Algorithms For Pairwise Protein Structure Alignment. PhD thesis, Vrije Universiteit, 01 2012.

[21] Faruck Morcos, Andrea Pagnani, Bryan Lunt, Arianna Bertolino, Debora S Marks, Chris Sander, Riccardo Zecchina, José N Onuchic, Terence Hwa, and Martin Weigt. Direct-coupling analysis of residue coevolution captures native contacts across many protein families. Proceedings of the National Academy of Sciences, 108(49):E1293–E1301, 2011.

[22] Carlo Baldassi, Marco Zamparo, Christoph Feinauer, Andrea Procaccini, Riccardo Zecchina, Martin Weigt, and Andrea Pagnani. Fast and accurate multivariate gaussian modeling of protein families: predicting residue contacts and protein-interaction partners. PloS one, 9(3):e92721, 2014.

[23] Magnus Ekeberg, Cecilia Lövkvist, Yueheng Lan, Martin Weigt, and Erik Aurell. Improved contact prediction in proteins: using pseudolikelihoods to infer potts models. Physical Review E, 87(1):012707, 2013.

[24] John P Barton, Eleonora De Leonardis, Alice Coucke, and Simona Cocco. Ace: adaptive cluster expansion for maximum entropy graphical model inference. Bioinformatics, 32(20):3089–3097, 2016.

[25] Daphne Koller and Nir Friedman. Probabilistic graphical models: principles and techniques. MIT press, 2009.

[26] Julian Besag. Statistical analysis of non-lattice data. Journal of the Royal Statistical Society: Series D (The Statistician), 24(3):179–195, 1975.

[27] Susann Vorberg. Bayesian Statistical Approach for Protein Residue-Residue Contact Prediction. PhD thesis, Ludwig-Maximilians-Universität, 2017.

[28] Stefan Seemayer, Markus Gruber, and Johannes Söding. Ccmpred—fast and precise prediction of protein residue–residue contacts from correlated mutations. Bioinformatics, 30(21):3128–3130, 2014.

[29] Milot Mirdita, Lars von den Driesch, Clovis Galiez, Maria J Martin, Johannes Söding, and Martin Steinegger. Uniclust databases of clustered and deeply annotated protein sequences and alignments. Nucleic acids research, 45(D1):D170–D176, 2017.

[30] Stefan Seemayer. Github ccmpred - frequently asked questions (faq). https://github.com/soedinglab/CCMpred/wiki/FAQ.

[31] J Michael Sauder, Jonathan W Arthur, and Roland L Dunbrack Jr. Large-scale comparison of protein sequence alignment algorithms with structure alignments. Proteins: Structure, Function, and Bioinformatics, 40(1):6–22, 2000.

[32] Julie D. Thompson, Frédéric Plewniak, and Olivier Poch. Balibase: a benchmark alignment database for the evaluation of multiple alignment programs. Bioinformatics (Oxford, England), 15(1):87–88, 1999.

[33] Robert C. Edgar. Qscore. http://www.drive5.com/qscore/.

[34] Richard H Lathrop. The protein threading problem with sequence amino acid interaction preferences is np-complete. Protein Engineering, Design and Selection, 7(9):1059–1068, 1994.

